# Dysconnectivity of the cerebellum and somatomotor network correlates with the severity of alogia in chronic schizophrenia

**DOI:** 10.1101/2024.04.16.589573

**Authors:** Wiktor Więcławski, Krzysztof Bielski, Martin Jani, Marek Binder, Przemysław Adamczyk

**Affiliations:** Institute of Psychology, Jagiellonian University, Krakow, Poland; Department of Psychiatry, Faculty of Medicine, Masaryk University and University Hospital Brno, Brno, Czech Republic

**Keywords:** schizophrenia, fMRI, somatomotor network, cerebellum, negative symptoms

## Abstract

Recent fMRI resting-state findings show aberrant functional connectivity within somatomotor network (SMN) in schizophrenia. Moreover, functional connectivity aberrations of the motor system are often reported to be related to the severity of psychotic symptoms. Thus, it is important to validate those findings and confirm their relationship with psychopathology. Therefore, we decided to take an entirely data-driven approach in our fMRI resting-state study of 30 chronic schizophrenia outpatients and 30 matched control subjects. We used independent component analysis (ICA), dual regression, and seed-based connectivity analysis. We found reduced functional connectivity within SMN in schizophrenia patients compared to controls and SMN hypoconnectivity with the cerebellum in schizophrenia patients. The latter is strongly correlated with the intensity of alogia, i.e. poverty of speech and reduction in spontaneous speech, demonstrated by patients. Our results are consistent with the recent knowledge about the role of the cerebellum in cognitive functioning and its abnormalities in psychiatric disorders, e.g. schizophrenia. In conclusion, the presented results, for the first time clearly showed the involvement of the cerebellum hypoconnectivity with SMN in the persistence and severity of alogia symptoms in schizophrenia.

## 1. Introduction

Alongside prominent positive and negative psychotic symptoms and cognitive impairment, most schizophrenia patients also experience a range of distressing motor symptoms (e.g., involuntary movement, catatonia, psychomotor slowing; Walther & Strik, 2012). Although motor symptoms have been considered primarily as side effects of antipsychotic treatment, they are also present in drug naïve and first-episode patients (Peralta et al., 2010). To investigate neural foundations of the above-mentioned phenomenon, resting-state functional magnetic resonance imaging (fMRI) has often been used, focusing on functional connectivity, i.e. measure of synchronization between remote neural assemblies (Fingelkurts et al., 2005). So far, evidence of resting-state functional connectivity changes compared to neurotypicals in various parts of the motor system has been found, including increased functional connectivity within basal ganglia (Duan et al., 2015), decreased functional connectivity of the cerebellum with multiple cortical regions (Andreasen & Pierson, 2008; Collin, 2011), and increased functional connectivity of primary motor cortex with the thalamus (T. Li et al., 2016)

Furthermore, decreased functional connectivity within somatomotor network (SMN; Keyvanfard et al., 2023) consisting of precentral (PRG) and postcentral (POG) gyri and supplementary motor area (SMA; Uddin et al., 2019) has been found in schizophrenia patients. What is more, the hypoconnectivity of the SMN and other resting state networks (Huang et al., 2020; Keyvanfard et al., 2023) is also present in this clinical population. These reported connectivity measures are often correlated with the severity of both positive and negative symptoms or impairment of cognitive performance (Bernard et al., 2017; Harikumar et al., 2023). Recent findings additionally suggest that dysconnectivity (a widely used term that can mean both hyper- and hypoconnectivity; Pettersson-Yeo et al., 2011) within that network can be a transdiagnostic marker of schizophrenia, bipolar disorder, and major depression (Huang et al., 2020; Magioncalda et al., 2020). All in all, current empirical research suggests that, alongside aberrations in other resting-state networks (e.g. hyperconnectivity within default mode network; Li et al., 2019; Mingoia et al., 2012), the motor system connectivity disturbances seem to play an important role in the clinical image of schizophrenia.

The primary objective of this study was to investigate between-group differences in functional connectivity of the SMN in chronic schizophrenia outpatients compared to neurotypicals. We acknowledge that schizophrenia is a heterogeneous disease, so to avoid arbitrarily choosing regions of interest (ROI) based on previous studies or atlases, we applied the data-driven approach to explore functional connectivity patterns. Furthermore, ROIs based on resting-state analysis proved to produce stronger activation in tasks when compared to ROIs based on meta-analyses or other single studies (Pamplona et al., 2020). For that reason, the probabilistic independent component analysis (ICA) was utilized to identify the SMN location and then, within-network connectivity patterns differences between patients and healthy controls were analyzed. Subsequently, seed-based functional connectivity analysis was performed using brain regions recognized in the previous step as seeds. Then, we looked for between-group differences in the functional connectivity of those regions with other brain areas.

The second aim was to investigate the relationships between aberrant brain connectivity with the severity of psychopathological symptoms. Therefore, we compared the strength of connectivity with individual results of psychiatric scales indicating levels of positive, negative, and disorganization symptoms. This comparison enables inferring the origins or implications of the observed correlations between brain networks and psychiatric symptomatology in these patients.

## 2. Methods

### 2.1 Subjects

The study involved two groups: 30 chronic schizophrenia patients (SCH) and 30 controls (CON) matched with respect to age and sex (Table I; the sample is described in detail elsewhere, Adamczyk et al., 2021). All participants signed the informed consent for the participation in the study. The study protocol was accepted by the Research Ethics Committee at the Institute of Psychology, Jagiellonian University, Krakow, Poland; following standards of the World Medical Association Declaration of Helsinki (2013).

**Table I.**
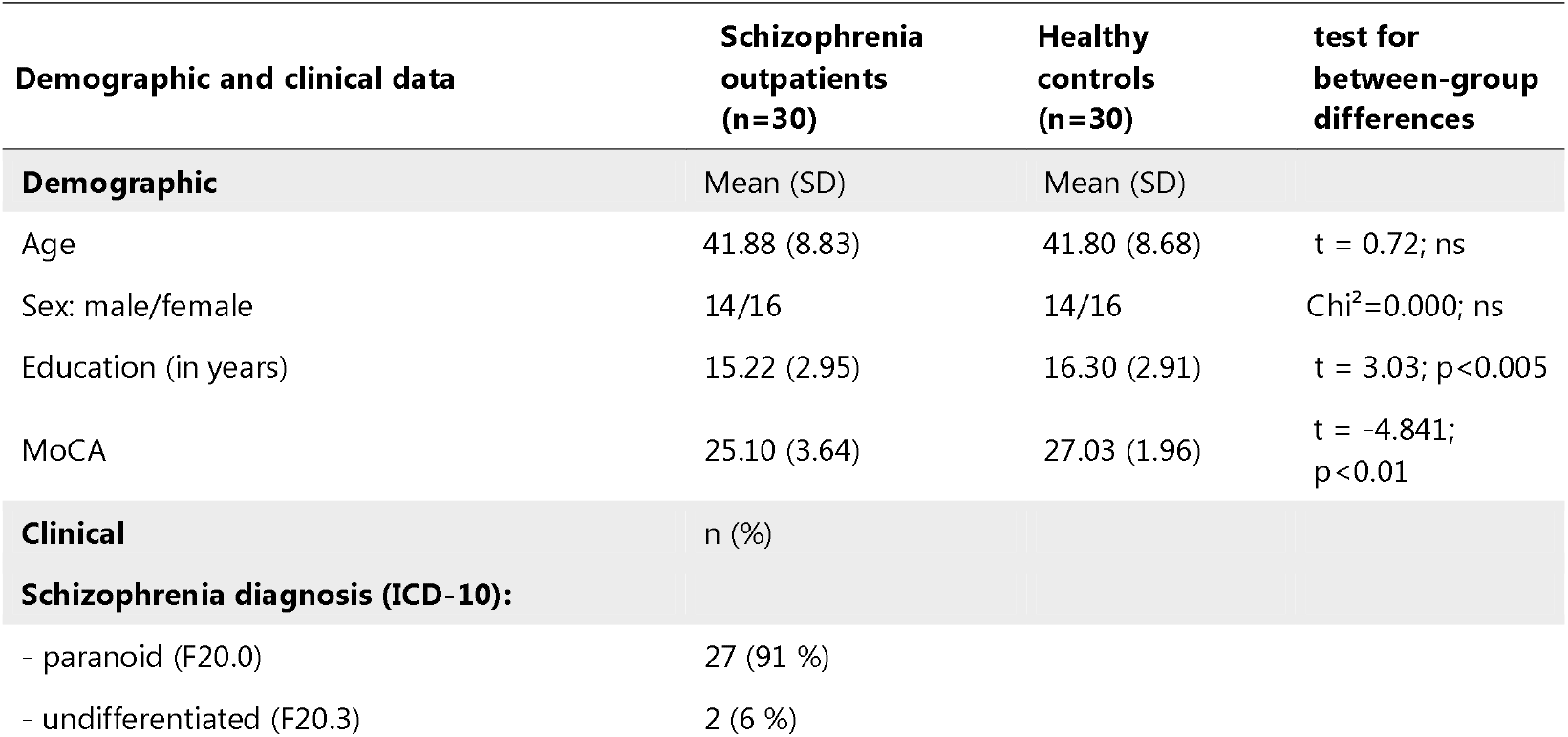
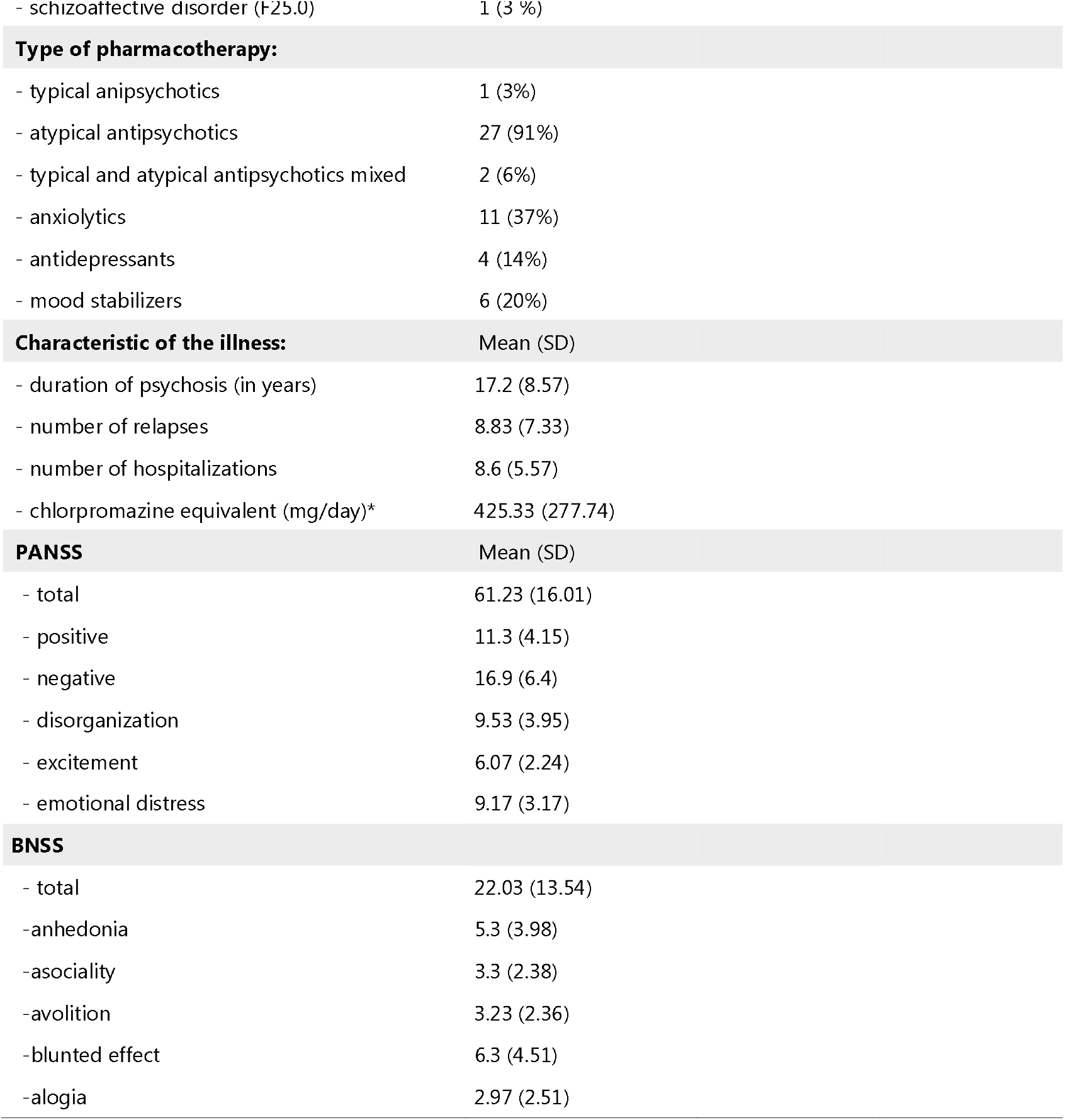
Demographic and clinical data.

Three subjects (two SCH and one CON) were removed from the further analysis due to excessive head movements in the magnetic resonance imaging (MRI) scanner.

Abbreviations: MoCA, Montreal Cognitive Assessment; PANSS, Positive and Negative Syndrome Scale; BNSS, Brief Negative Symptom Scale; ICD-10, International Classification of Diseases, Tenth Revision. Subjects’ demographics and clinical data were presented as mean (SD) for quantitative data and n (%) for the nominal variable. The significance level in all statistical analyses equaled alpha = 0.05. Typical antipsychotics: flupentixol, haloperidol, promazine; atypical antipsychotics: amisulpride, clozapine, olanzapine, risperidone, sulpiride, quetiapine; aripiprazole; antidepressants: escitalopram, paroxetine; anxiolytics: hydroxyzine; mood stabilizers: carbamazepine, lithium, valproic acid.

### 2.2 Severity of psychopathology symptoms assessment

The severity of psychopathology symptoms in the schizophrenia patients was measured before MRI scanning. Montreal Cognitive Assessment (MOCA; Nasreddine et al., 2005) Positive and Negative Syndrome Scale (PANSS; Kay et al., 1987; Vandergaag, Cuijpers, et al., 2006) and Brief Negative Symptoms Scale (BNSS; (Kirkpatrick et al., 2011) were administered by an experienced psychologist and psychiatrist to measure respectively, the cognitive performance and severity of psychopathology. The five-factor PANSS (Vandergaag, Hoffman, et al., 2006) subscales measures: a) positive (e.g., hallucinations, delusions, paranoid), b) negative (e.g., blunted affect, social withdrawal, motor retardation), c) disorganization (cognitive deteriorations e.g., conceptual disorganization, diminished abstract thinking, poor attention), d) excitement (e.g., excitement, hostility, impulse control) and e) depression/anxiety (e.g., guilt, tension, depression, and anxiety) symptoms. The five-factor BNSS (Ahmed et al., 2019) provides more precise information on the five core negative symptoms, i.e.: a) alogia (poverty of speech), b) anhedonia (lack of ability to experience pleasure), c) avolition (low energy and motivation, diminished interest in usual activities), d) asociality (e.g. social withdrawal, disrupted social interactions and avoidance of interpersonal connections) and e) blunted affect (i.e., diminished response to emotional stimuli).

### 2.3 MRI resting-state procedure

Before entering MRI scanner, participants were instructed on the safety issues in MRI and asked to relax, lie still, refrain from any head movement, and focus their gaze on the fixation cross during 10 minutes of the protocol.

### 2.4 MRI image acquisition

MRI data acquisition was performed with a 3T scanner (Magnetom Skyra, Siemens) at Malopolska Centre of Biotechnology, Krakow, Poland. The participants’ heads were immobilized with pillows. The resting-state functional images were acquired in the interleaved fashion. The scan parameters were as follows: TR = 2000 ms, TE = 27 ms, FA = 75, matrix size = 64x64, field of view = 1344 mm × 1344 mm, number of slices = 35, voxel size = 3x3x3 mm, and slice thickness = 3 mm (inter-slice gap = 3.3 mm). A total of 300 volumes were acquired. Anatomical images were acquired using the T1 MPRAGE sequence (sagittal slices: 1x1x1.1 mm3 voxel size; TR=1800 ms, TE=2.37 ms).

### 2.5 Data preprocessing

Data were preprocessed using CONN software version 22a (Nieto-Castanon & Whitfield-Gabrieli, 2022). Functional and anatomical data were preprocessed using a flexible preprocessing pipeline (Nieto-Castanon, 2020): 1) removal of 30 initial scans in each subjects’ functional run to account for time when written instruction was displayed and a short period of adaptation to the scanner for the subjects, 2) realignment with susceptibility distortion correction using fieldmaps (realign & unwarp procedure (Andersson et al., 2001) integrating fieldmaps for susceptibility distortion correction; all scans were coregistered to a reference image (first scan of the first session) using a least squares approach and a 6 parameter (rigid body) transformation), 3) slice timing correction, 4) outlier detection (ART toolbox - framewise displacement above 0.5 mm or global BOLD signal changes above 3 standard deviations), 5) direct segmentation (Tissue Probability Maps), 6) MNI-space normalization, and 7) smoothing with a Gaussian kernel of 6 mm FWHM. In addition, functional data were denoised using a standard denoising pipeline (Nieto-Castanon, 2020) including the regression of potential confounding effects characterized by white matter timeseries (5 CompCor noise components), CSF timeseries (5 CompCor noise components), motion parameters and their first order derivatives (12 factors), outlier scans (below 168 factors), session effects and their first order derivatives (2 factors), and linear trends (2 factors) within each functional run, followed by bandpass frequency filtering of the BOLD timeseries between 0.008 Hz and 0.09 Hz. CompCor noise components within white matter and CSF were estimated by computing the average BOLD signal as well as the largest principal components orthogonal to the BOLD average, motion parameters, and outlier scans within each subject’s eroded segmentation masks.

### 2.6 Data analysis

#### 2.6.1 Defining SMN and other networks

Probabilistic ICA (Beckmann & Smith, 2004) as implemented in MELODIC (Multivariate Exploratory Linear Decomposition into Independent Components) Version 3.15, part of FSL (FMRIB’s Software Library, www.fmrib.ox.ac.uk/fsl) was performed to identify the general location and temporal characteristics of resting-state brain networks. The number of components was estimated to be 14 using the minimum description length criterium (Rissanen, 1978).

#### 2.6.2 Recognizing regions of interest

The set of spatial maps from the group-average analysis was used to generate subject-specific versions of the spatial maps, and associated timeseries, using dual regression (Nickerson et al., 2017). We then tested between-group differences using FSL’s randomise permutation-testing tool (Winkler et al., 2014) with Family Wise Error (FWE) correction on threshold-free cluster enhancement at alpha set to 0.05.

#### 2.6.3. Seed-based connectivity

Regions that differed significantly between groups in the previous step were taken as seeds for further analysis in the CONN software. Seed-based connectivity maps (SBC) were estimated using the Fisher-transformed bivariate correlation coefficients from a weighted general linear model (weighted-GLM; Nieto-Castanon, 2020). In each SBC map, coefficients were estimated separately for each seed and target voxels, modelling the association between their BOLD signal time-series. In order to compensate for possible transient magnetization effects at the beginning of each run, individual scans were weighted by a step function convolved with an SPM canonical hemodynamic response function and rectified.

Group-level analyses were performed using a General Linear Model (GLM). Voxel-level hypotheses were evaluated using multivariate parametric statistics with random-effects across subjects and sample covariance estimation across multiple measurements. Inferences were performed at the level of individual clusters. Cluster-level inferences were based on parametric statistics from Gaussian Random Field theory (Worsley et al., 1996). Results were thresholded using a combination of a cluster-forming *P* < 0.001 voxel-level threshold, and a False Discovery Rate (FDR) *P* < 0.05 cluster-size threshold (Chumbley et al., 2010).

#### 2.6.4 Relationship of connectivity and psychiatric measures

Parameter estimates from dual regression and connectivity values between a seed and significant cluster were extracted from each subject and imported to JASP (Version 0.18.3; JASP Team, 2024) to calculate Spearman correlation coefficients with PANSS and BNSS subscales. We decided to correct for a number of clinical scales that we tested for PANSS and BNSS factorial models separately (7 comparisons for BNSS and 6 comparisons for PANSS).

Effect sizes for mean cluster connectivity between-group differences were estimated by calculating rank-biserial correlation (r_rb_) as variables’ distributions deviated from normal.

## 3. Results

### 3.1. Recognizing SMN

All ICA group components were inspected by experienced neuroscientists (M.B., P.A.) One component closely resembled SMN. It displayed the strongest activation in PRG, POG, and SMA regions. We used the Personode software (Pamplona et al., 2020) to determine the similarity of networks we obtained with the predefined SMN template. The correlation between our IC components and the template was the highest for the component we identified as SMN (58%).

### 3.2. Recognizing regions of interest for seed-based connectivity analysis

Between-group comparisons revealed significantly decreased functional connectivity of the left SMA within the SMN in SCH (T = 4.64, *p* = 0.013, r_rb_ = 0.98, cluster size = 172; peak MNI coordinates = -2 -18 44; Fig. 1A). This one cluster was selected for performing the seed-based correlations.

**Fig. 1.**
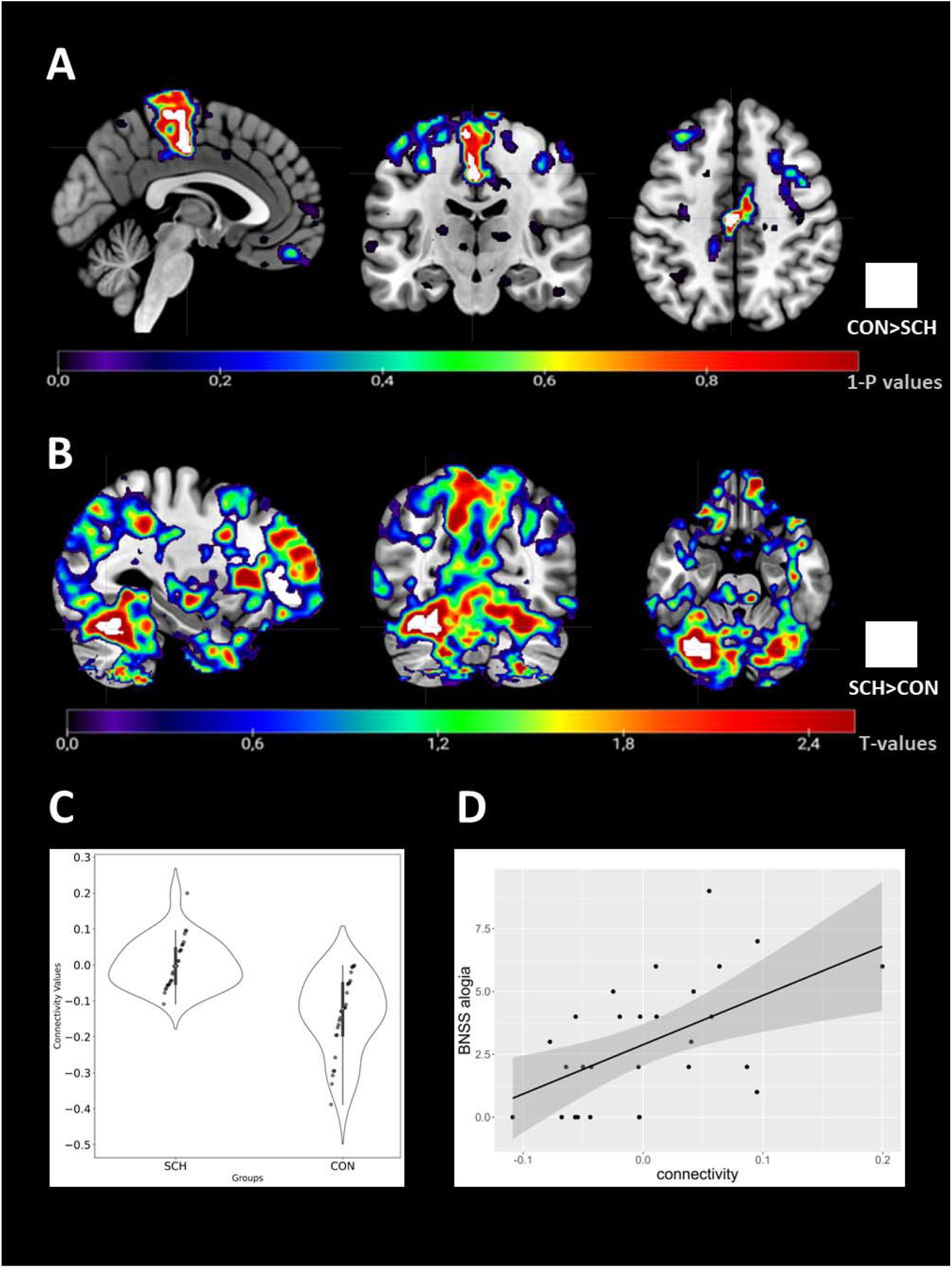
Abnormal SMN-cerebellum connectivity and its relationship with BNSS alogia in schizophrenia. A) Between-group differences in SMN connectivity. FWE-corrected results at contrast CON > SCH with no threshold. Slices presented at MNI: x=-2; y= -18; z=44. A statistically significant cluster at *p* < 0.05 is highlighted in white. The color bar represents 1 – *p* values. B) Between-group differences in SMN - cerebellum seed-based connectivity at contrast SCH > CON. Results with no threshold. A statistically significant cluster at *p* < 0.05 within grey matter is highlighted in white. The color bar represents T values. Slices presented at MNI: x=-30; y= -68; z= -26. C) SMA-cerebellar lobule VI connectivity values violin plot in both groups. SCH connectivity clearly oscillates around zero while CON subjects show tendency toward negative functional connectivity D) Scatterplot of SMA-cerebellar lobule VI connectivity values and BNSS alogia scores with density distribution of both variables. Abbreviations: SCH – chronic schizophrenia outpatients, CON – controls, BNSS – Brief Negative Symptoms Scale.

### 3.3. Seed-based connectivity patterns

The left SMA exhibited negative connectivity with the left cerebellar lobule VI (Table II; Fig. 1B) in the CON group (*r* = -0.17) and near-zero connectivity in the SCH group (*r* = 0.01). The difference between the two groups was statistically significant (*p*<0.001; Fig. 1C).

**Table II.** Conn seed-based connectivity results.

| Seed area | Target area | Cluster size | T | Peak voxel significance | Cluster significance | $r_{rb}$ (95% CI) | Peak voxel coordinates (x, y, z) |
| --- | --- | --- | --- | --- | --- | --- | --- |
| Supplementary motor area | Cerebellar lobule VI | 248 | 6.25 | $p = 0.246$ | $p < 0.001$ | 0.76 (0.61-0.86) | -30 -68 -26 |

Statistically significant FDR-corrected results of between group contrast. Only clusters within grey matter are reported. Abbreviations: CI – confidence intervals, $r_{rb}$ – rank-biserial correlation

**Table III.** SMA-cerebellar lobule VI mean cluster connectivity and clinical scales correlation analysis results.

| Clinical (sub)scale | Spearman's rho (95% CI) | p | p FDR |
| --- | --- | --- | --- |
| PANSS total | 0.30 (-0.11-0.65) | 0.139 | 0.348 |
| PANSS positive | 0.14 (-0.28-0.55) | 0.509 | 0.611 |
| PANSS negative | 0.41 (0.00-0.72) | **0.039\*** | 0.234 |
| PANSS disorganization | 0.28 (-0.12-0.61) | 0.174 | 0.348 |
| PANSS excitement | 0.17 (-0.23-0.55) | 0.397 | 0.596 |
| PANSS emotional distress | 0.03 (-0.39-0.43) | 0.905 | 0.905 |
| BNSS total | 0.45 (0.04-0.74) | **0.023\*** | 0.053 |
| BNSS anhedonia | 0.16 (-0.27-0.57) | 0.437 | 0.437 |
| BNSS distress | 0.43 (0.06-0.72) | **0.030\*** | 0.053 |
| BNSS asociality | 0.34 (-0.60-0.64) | 0.092 | 0.107 |
| BNSS avolition | 0.36 (-0.04-0.67) | 0.074 | 0.104 |
| BNSS blunted affect | 0.43 (-0.01-0.75) | **0.028\*** | 0.053 |
| BNSS alogia | 0.53 (0.14-0.78) | **0.006\*\*** | **0.042\*** |

### 3.4 Relationship between connectivity and the clinical symptoms measures

When controlling for illness duration and chlorpromazine equivalent of medication SMA connectivity within SMN from dual regression analysis did not correlate significantly with clinical scales. Meanwhile, mean cluster connectivity values between the left SMA and left cerebellar lobule VI from seed-based analysis were significantly related to clinical scores in SCH (Table IV). SMA-VI lobule connectivity score correlated with a total score of BNSS, BNSS alogia, blunted affect, distress, and PANSS negative symptoms scale. Finally, Only BNSS alogia survived FDR correction for multiple comparisons (Fig. 1D).

Spearman’s rho partial correlation coefficients with bootstrapped (10,000 samples) confidence intervals are shown. Illness duration and chlorpromazine equivalent of medication are controlled. Abbreviations: CI – confidence intervals, FDR – false discovery rate. ^*^ *p* < 0.05; ^**^ *p* < 0.01

## 4. Discussion

In the present paper, we investigated intrinsic functional connectivity of SMN, using the whole-brain data-driven approach. By applying the analytical pipeline involving ICA decomposition, dual regression and seed-based correlations analysis, we found hypoconnectivity within the SMN and its dysconnectivity with the cerebellum in a sample of patients with schizophrenia when compared with the neurotypical control group.

More specifically, we found that aberrant SMN-cerebellum connectivity (i.e., between the medial SMA and the left cerebellar lobule VI) was correlated significantly with the BNSS alogia subscale. Interestingly, it may indicate that disruption of SMN-cerebellum connectivity may be recognized as related to motor-based problems, resulting in enhancement of manifestation of specific negative symptoms, such as pronounced reduction in spontaneous speech, a narrowing of the range of speech, and deficiencies in speech content.

Our results are consistent with the current knowledge on the involvement of the cerebellum in language processing (De Smet et al., 2007; Mariën & Beaton, 2014; Zhang et al., 2023). The left VI cerebellar lobule that we report is known to be functionally connected to SMN at rest and involved in motor tasks (Guell et al., 2018). The general role of the cerebellum in cognitive processing and its abnormal functioning in schizophrenia has been an influential idea in the past decade. The cerebellum is mainly responsible for the prediction of sensorimotor input and the correction of errors that arise during the planning and execution of movement (Albus, 1971; Blakemore et al., 2000; Marr, 1969). In the recent models of cerebellar function, predicting and detecting errors is extended beyond the motor domain to the cognitive domain, e.g. predicting the next word in a sentence (D’Mello et al., 2020; Leiner et al., 1991). We know that the cerebellum contributes to verbal fluency (Molinari & Leggio, 2016) which is also commonly impaired in schizophrenia (Gourovitch et al., 1996). Thus, the dysconnectivity of the cerebellum may correspond to abnormal adaptive feedback to the cortex concerning pattern changes and error information.

Therefore, this error monitoring process seems to be extremely important in various language functions, which is in line with the direct cerebellar contribution to language hypothesis (Daum & Ackermann, 1995; De Smet et al., 2007). The reported SMN-cerebellum dysconnectivity requires future experimental studies, which might reveal the importance of such a phenomenon and its impact on alogia manifestation and persistence. Noteworthy, in light of our findings, we may consider alogia as a severe motor-based impairment of verbal fluency.

Furthermore, our results are consistent with previous findings on the dysconnectivity of the cerebellum with cortical regions and altered connectivity within cerebellum, both linked to the severity of negative symptoms in previous studies (Choi et al., 2023; Kim et al., 2014). Importantly, it has been shown that stimulation of the cerebellum with transcranial magnetic stimulation (TMS) in patients with altered cerebellar-prefrontal network connectivity was reported to cause a reduction in negative symptoms (Brady et al., 2019; Hua et al., 2022). One of the reasons why TMS therapy is effective in reducing negative symptoms may be the establishment of synchronization between SMN and the cerebellum.

Finally, it is worth mentioning that, most previous studies in their correlation analysis used just one general scale to measure negative symptoms (PANSS), so they do not provide us with more insight into aetiology of its specific symptomatology. Therefore, our work provides a novel insight into the specificity of negative symptoms subscales and their relationships with altered brain connectivity, i.e. SMA-cerebellum and alogia. The small sample size is a main limitation of this study thus its result should be interpreted as preliminary, and need further corroboration with a larger patient sample. Therefore, future studies on bigger samples with specific clinical assessment are needed to resolve the question of the role of SMA-cerebellum connectivity and future neurotherapeutic intervention (e.g. TMS, tDCS) on its effectiveness in the reduction of alogia in schizophrenia individuals.

## Conclusions

Our study highlights the hypoconnectivity within the SMN and SMN dysconnectivity with the cerebellum. The latter connection is linked to alogia. The involvement of the cerebellum in language processing and its aberrant connectivity with the SMN underscore its potential role in the clinical manifestation of alogia. Further research on interventions targeting cerebellar connectivity, such as TMS, holds promise for alleviating negative symptoms in schizophrenia and revealing in more detail neural underpinnings of alogia.

## Acknowledgments

The study was supported by the National Science Centre Poland under grant no. 2016/23/B/HS6/00286 and 2021/41/B/HS6/02967. We cordially thank Olga Dudzińska for her help in data acquisition, all the subjects for participation, and psychiatrists for their assessment.

## Authors Contribution (CRediT)

WW: Conceptualization, Formal Analysis, Methodology, Visualization, Writing – original draft; KB: formal analysis, methodology, writing – review and editing; MJ: Investigation, Writing – Review & Editing; MB: Writing - Review & Editing, Formal analysis; PA: Conceptualization, Methodology, Investigation, Project Administration, Funding Acquisition, Resources, Writing - Review & Editing;

## Ethical approval

All procedures performed in studies involving human participants were in accordance with the ethical standards of the institutional national research committee (The Research Ethics Committee at the Institute of Psychology, Jagiellonian University KE/02/082017) and with the 2013 World Medical Association Declaration of Helsinki (2013).

## Informed consent

Informed consent was obtained from all individual participants included in the study.

